# Intradermal Administration of Rescue Dose of Antiemetics in Chemotherapy-induced Nausea and Vomiting

**DOI:** 10.1101/2023.09.27.559650

**Authors:** Minami Maekawa-Matsuura, Natsuko Shimizu, Yuri Sakamaki, Eri Oya, Sakiko Shimizu, Yoichiro Iwase

**Author notes:** Corresponding author at: 1500 Inokuchi, Nakai-machi, Ashigarakami-gun, Kanagawa 259-0151, Japan. E-mail address (Y. Iwase).

## Abstract

Antiemetic therapy is provided to patients with cancer to prevent chemotherapy-induced nausea and vomiting (CINV), based on the emetic risk of anticancer agents. However, breakthrough CINV can occur despite appropriate antiemetic prophylaxis. It is preferable to administer the rescue dose of antiemetics in breakthrough CINV that occur at home, in a form that is immediately effective and that can be easily self-administered by the patient. In the present study, we examined and compared the blood pharmacokinetics of subcutaneous (SC) injection, a preferred route of self-injection, and intradermal (ID) injection, a less invasive route, with an aim of developing simple self-injectable rescue antiemetics. A single administration of four kinds of antiemetics via SC and ID routes in rats resulted in lower *T*_max_ values compared with previously reported oral administration, indicating that the administration of antiemetics via ID and SC routes may be effective in providing faster antiemetic effects and to alleviate patient symptoms.

## Introduction

Chemotherapy-induced nausea and vomiting (CINV) is an adverse effect of cancer therapy that reduces the quality of life of the patients. CINV is classified into the following five categories: acute, delayed, breakthrough, anticipatory, and refractory. Breakthrough CINV is defined as nausea and/or vomiting occurring within 5 days of chemotherapy despite guideline-based antiemetic prophylaxis (1). Breakthrough CINV is difficult to control and may result in interruption or discontinuation of cancer therapy (2). When breakthrough CINV occurs, it is recommended to use scheduled doses of multiple antiemetics with different mechanisms of action or to switch to a 5-HT_3_ receptor antagonist that is different from the one used for prophylaxis. In a prospective observational study of adult patients receiving their first highly emetogenic or moderately emetogenic chemotherapy (HEC or MEC) (2001–2002, the US and Europe), at least 35% of all patients experienced acute nausea and 13% experienced acute vomiting after antiemetic prophylaxis. Additionally, delayed nausea and vomiting were observed in 60% and 50% of HEC patients and 52% and 28% of MEC patients, respectively (3). According to a prospective, multicenter, and observational study (2011– 2012, Japan), 943 (49%) of 1910 patients who received HEC or MEC developed breakthrough CINV, and 412 out of the 943 patients received rescue medication (4). In a study of patients receiving HEC or MEC, a significant improvement in complete response (no vomiting and no use of rescue medication) during the delayed and overall period was noted in patients treated with olanzapine (an atypical antipsychotic) as compared with the control group (5). Based on these results, the 2016 revised guidelines of the Multinational Association of Supportive Care in Cancer (MASCC) and European Society of Clinical Oncology (ESMO) recommended the use of 10 mg oral olanzapine for 3 days for breakthrough CINV (6). Although the prevention of vomiting is fairly managed by the inclusion of olanzapine in the treatment of breakthrough CINV, nausea requires additional medical attention (6). Further research is needed for CINV that occurs at home because there are no clinical studies focusing on effective therapies for delayed CINV (7). Keeping additional doses of rescue medication accessible for treating breakthrough CINV can be beneficial in timely management of nausea in patients. Furthermore, there is a need for an administration route that is comfortable to patients and has rapid absorption. (8, 9). However, the most employed administration routes for antiemetics in chemotherapy regimens are the intravenous (IV) and per os (PO) routes (10). IV administration has a faster onset of action, but it is difficult for patients to intravenously self-administer at home. Intramuscular (IM) injections, which have pharmacokinetics similar to those of IV injections, enables a rapid onset of action; however, it carries a risk of vascular or neurological impairment and is accompanied by pain and fear (11, 12). Antiemetic administration via the subcutaneous (SC) route, which is commonly used to administer self-injectable formulations, has been considered as an alternative administration route for outpatients who are unable to tolerate oral medications due to vomiting or other reasons, or when IV administration is not possible (13, 14). However, in the case of sumatriptan, a drug for treating cluster headache, some patients are reported to prefer nasal administration due to its convenience, despite the low efficacy of nasal administration compared to SC administration (15). In comparison to oral sumatriptan tablets, 47% of patients preferred tablets due to its convenience over SC self-injection, which also showed higher efficacy than tablets (16). Nevertheless, many patients receiving chemotherapy are unable to swallow or ingest antiemetic tablets because of vomiting, oral mucositis, or the presence of oral feeding tubes, and require another treatment options (17).

In SC administration, it has been reported that injection comfort can be improved by selecting smaller needles seizes (both gauge and length) (18–21). Therefore, we focused on intradermal (ID) injection, which targets a shallower area compared with the target area of SC injection and may increase convenience and reduce pain and apprehension compared with SC injection.

In the administration of insulin and low-molecular-weight headache drugs, ID administration is reported to be a useful administration route that can provide a time to maximum plasma concentration after drug administration (*T*_max_) that is comparable to or shorter than that of SC administration (22–24). Despite their interesting pharmacokinetics, ID injections have not been widely used clinically due to complexities of the Mantoux technique (25), which has traditionally been used for ID injection (26, 27). The Mantoux technique has not been applied to the injection of general drug solutions, because the insertion angles are low (5–15 degrees) and it is difficult to accurately administer drug solution intradermally due to the risk of drug solution transferring to the SC area or leaking from the skin surface (26, 27). As a remedy, various ID injection devices have been developed (28). However, as far as we have been able to find, there are no approved ID self-administration devices for liquid medications.

In the present study, we focused on our previously developed Immucise Intradermal Injection System (Immucise, Terumo Corporation (needle gauge, 33 G; needle length, 1.15 mm; bevel length, 0.6 mm)) (Figure 1) (29). Additionally, we had developed an improved version of the Immucise device for rats in a previous study (ID injection system for rats (IDS for rats) (needle gauge, 34 G; needle length, 0.5 mm; bevel length, 0.2 mm)) (30). In this study, we examined the possibility of easy-to-use, self-injectable rescue antiemetics at home to be administered subcutaneously or intradermally, by evaluating the plasma drug concentration after administering single doses of four types of antiemetic drugs to rats by SC injection and ID injection using the DS for rats.

**Figure 1.**
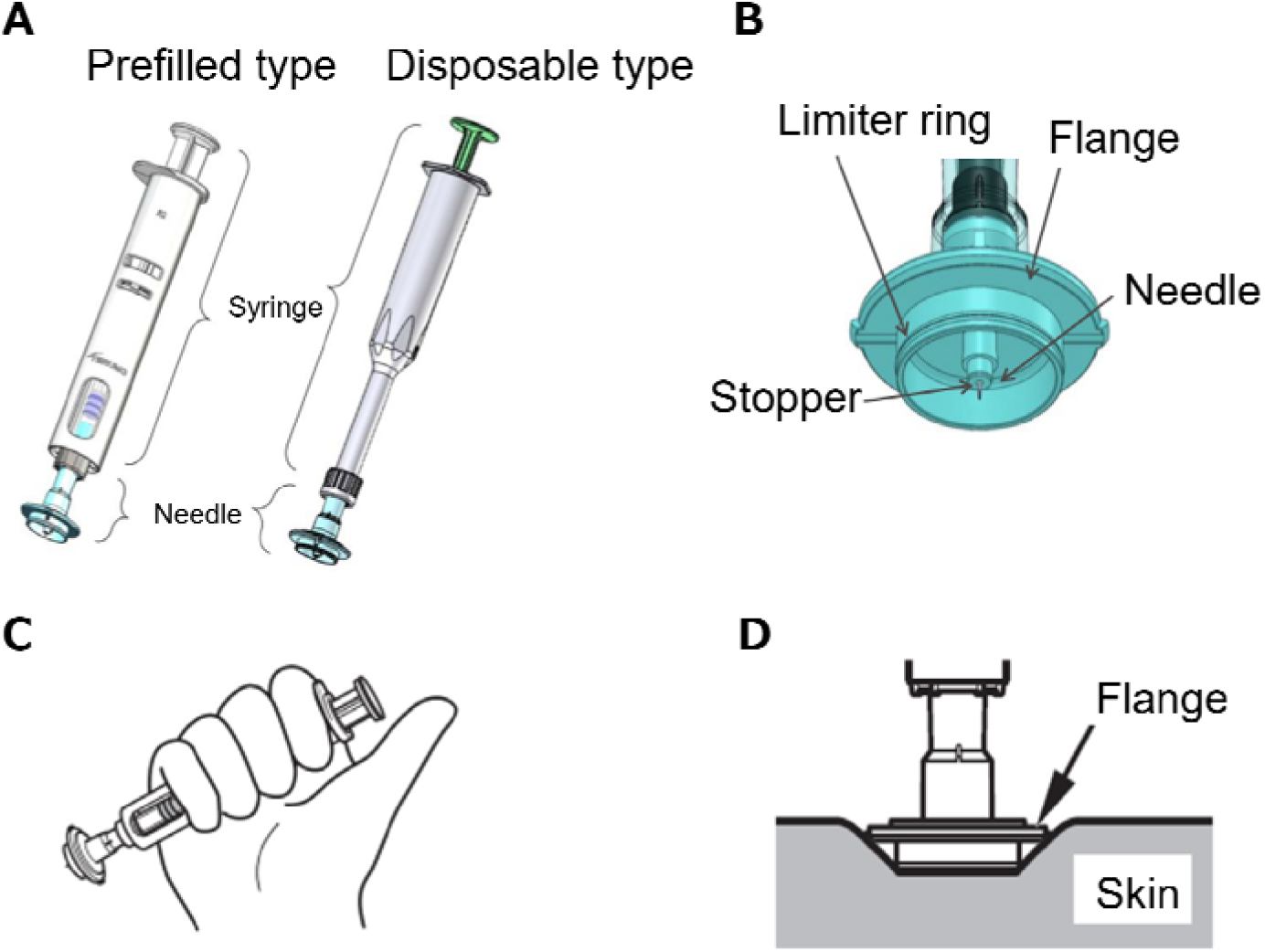
Immucise Intradermal Injection System. A: Two types of Immucise. B: Structure of Immucis needle. C: How to grip Immucise. D: Puncture image of Immucise vertically against the skin.

## Materials and Methods

### Animals

Male Sprague-Dawley (SD) rats (210–270 g, 7-week-old, Jackson Laboratory Japan, Kanagawa, Japan) were used in the study after a 6–7 day acclimation period. The rats were maintained in a temperature- and humidity-controlled environment and a 12-h light/dark cycle and had free access to food and water. Ethical approval for the present study was obtained from the Animal Experiment Committee of Terumo Corporation.

### Preparation of Administration Solution and Property Evaluation

Azasetron solution was prepared by dissolving 22.08 mg of azasetron hydrochloride (MedChem Express, NJ, USA) in 250 μL of water for injection and adding 250 μL of PBS (Nacalai Tesque, Kyoto, Japan). Olanzapine solution was prepared by dissolving Zyprexa for intramuscular injection (11.0 mg as olanzapine per vial, Eli Lilly Japan, Hyogo, Japan) in 0.525 mL of water for injection. Domperidone solution was prepared by dissolving 7.52 mg of domperidone (Tokyo Chemical Industry, Tokyo, Japan) in 9 μL of acetic acid (Kanto Chemical, Tokyo, Japan) and adding 738 μL of 0.01 mol/L acetic acid buffer solution (prepared by 0.1 mol/L acetic acid buffer solution (Nacalai Tesque, Kyoto, Japan) and water for injection) and 5 μL of 10 mol/L sodium hydroxide solution (Nacalai Tesque). Chlorpromazine solution was prepared using Contomin Intramuscular Injection (Mitsubishi Tanabe Pharma, Tokyo, Japan). The final concentrations of each drug solution were as follows: Azasetron solution, 40 mg/mL; olanzapine solution, 20 mg/mL; domperidone solution, 10 mg/mL; and chlorpromazine solution, 9 mg/mL. The pH was measured with a pH meter (LAQUA act D-72, Horiba, Kyoto, Japan). Osmotic pressure was measured with a single-sample osmometer (Instruments 3250 Osmometer, Advanced Instruments, Norwood, MA, USA). The molecular weight, logP value, pH, and osmolarity of each drug solution are shown in Table 1.

**Table 1.**
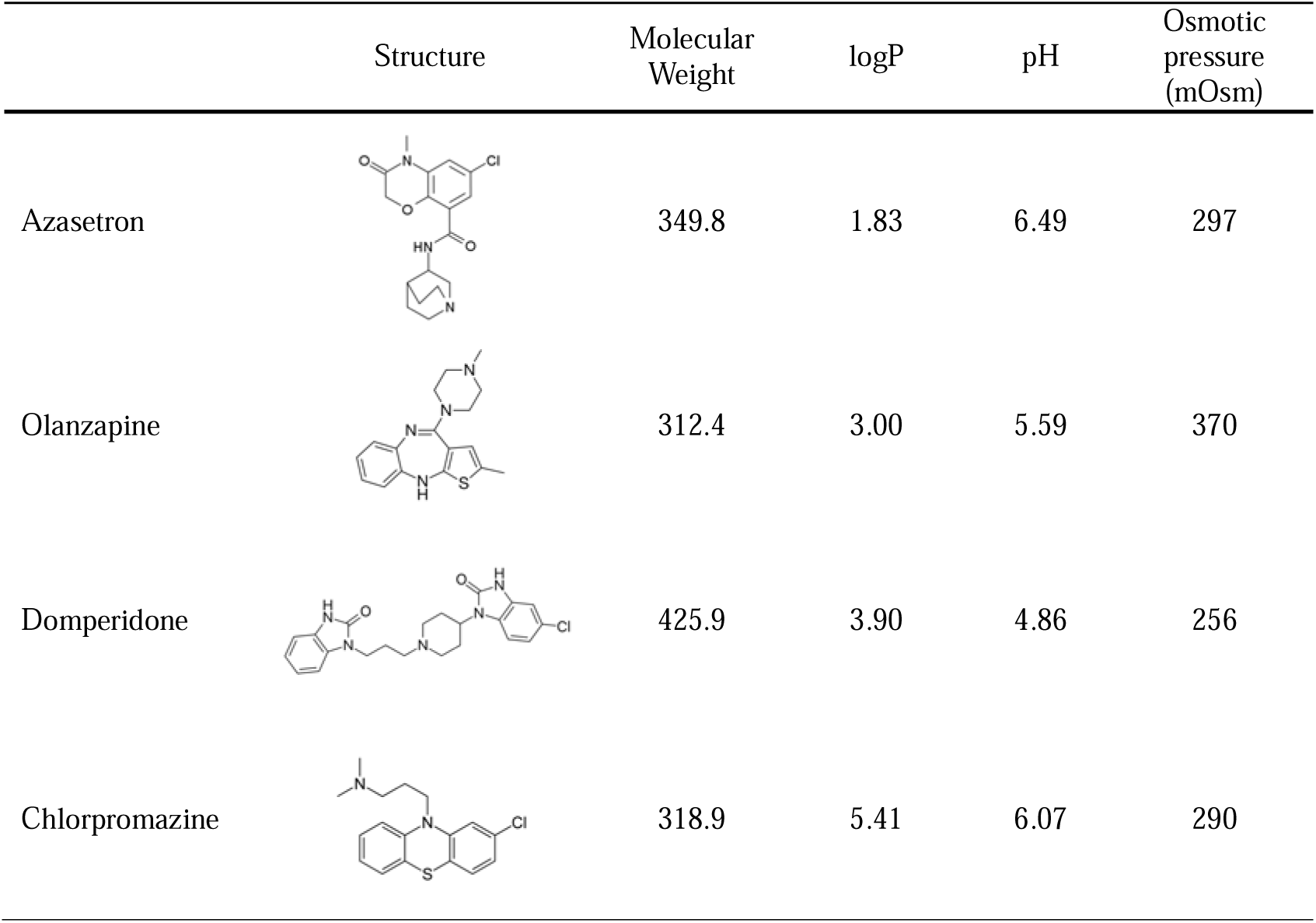
Characteristics of drugs and their solutions.

|  | Structure | Molecular Weight | logP | pH | Osmotic pressure (mOsm) |
| --- | --- | --- | --- | --- | --- |
| Azasetron |  | 349.8 | 1.83 | 6.49 | 297 |
| Olanzapine |  | 312.4 | 3.00 | 5.59 | 370 |
| Domperidone |  | 425.9 | 3.90 | 4.86 | 256 |
| Chlorpromazine |  | 318.9 | 5.41 | 6.07 | 290 |

### Drug Administration and Blood Sample Collection

Each solution was injected into the back of rats anesthetized by inhalation of isoflurane (Pfizer Japan, Tokyo, Japan). The volume of solution to be administered was 50 μL or less, and drug doses per kg of body weight of rats were: azasetron 5.5 mg/kg; olanzapine 2.5 mg/kg; domperidone; 1.4 mg/kg and chlorpromazine 1.2 mg/kg.

Azasetron and olanzapine were administered to three rats each, and domperidone and chlorpromazine were administered to five rats each.

For SC administration, a micro syringe with a 25 G fixed needle (Trajan Scientific Japan, Kanagawa, Japan) or a Hamilton micro syringe (Hamilton, Reno, NV, USA) connected to a 23 G injection needle (Terumo, Tokyo, Japan) was used. For ID administration, IDS for rats was used (30). The administration site was shaved in advance, and skin thickness was measured using an ultrasound diagnostic device (LOGIQ e Premium Pro, L10-22-RS probe, GE Healthcare Japan, Tokyo, Japan) and an ultrasound gel (LOGIQLEAN bottle hard type, GE Healthcare Japan) (30). Successful administration was defined as the absence of leakage of the injection solution and a wheal diameter of at least 3 mm confirmed visually after administration (30).

Blood was collected at 8 time points in total (before administration and at 5, 10, 15, 30, 45 min, 1 h, and 2 h after administration) for azasetron and olanzapine at 10 time points (before administration and at 5, 10, 15, 30, and 45 min and 1, 2, 4, and 24 h after administration) for domperidone; and at 14 time points (before administration and at 5, 10, 15, 30, and 45 minutes and 1, 2, 3, 4, 5, 6, 24, and 48 h after administration) for chlorpromazine. In this study, 0.2–0.3 mL of blood was collected from the jugular vein per blood draw. Heparin sodium (Mochida Pharmaceutical, Tokyo, Japan) was used as an anticoagulant. Plasma samples were obtained by centrifugation and stored frozen until further analysis.

### Standards and Internal standards

As the reference material and internal standard (IS), azasetron hydrochloride (Toronto Research Chemicals, Toronto, Canada) and azasetron-d3 hydrochloride (Toronto Research Chemicals) were used for azasetron; olanzapine (Tokyo Chemical Industry) and olanzapine-d8 (CAYMAN CHEMICAL) for olanzapine; domperidone (Tokyo Chemical Industry) and domperidone-d6 (Toronto Research Chemicals) for domperidone; and chlorpromazine hydrochloride (FUJIFILM Wako Pure Chemical, Osaka, Japan) and chlorpromazine-d6 hydrochloride (Toronto Research Chemicals) were used for chlorpromazine. Each IS solution was prepared using water, methanol, ethanol, or a mixture of these materials to obtain the concentrations of the samples described in the subsequent section.

### Sample Preparation

Azasetron-containing plasma samples were purified by deproteinization. Ten microliters of a mixture of water and methanol (1:1 (v/v)) and 10 μL of the IS solution (0.2 μg/mL) were added to 10 μL of each plasma sample and stirred. Subsequently, 100 μL of a mixture of acetonitrile and formic acid (1000:1 (v/v)) was added to each mixture and stirred. After centrifugation, 50 μL of the organic solvent layer was collected from each mixture, 200 μL of a mixture of water and formic acid (1000:1 (v/v)) was added, and the resultant solutions after stirring were used as measurement samples. Olanzapine- containing plasma was purified by liquid-liquid extraction. Thereafter, 10 μL of ethanol and the IS solution (0.4 μg/mL) were added to 10 μL of each plasma sample and stirred. Thereafter, 100 μL of 10 mmol/L ammonium formate was added to each mixture and stirred; 600 μL of t-butyl methyl ether was then added and the mixtures were stirred.

After centrifugation, 200 μL of the organic solvent layer was collected from each mixture and dried under a nitrogen stream at 40 degrees Celsius. Methanol (200 μL) was added to each residue and stirred. Additionally, 200 μL of water was added, and the resultant solutions after stirring were used as measurement samples. Domperidone-containing plasma samples were purified using deproteinization. Ten microliters of methanol and 10 μL of the IS solution (0.05 μg/mL) were added to 30 μL of each plasma sample and stirred. Next, 150 μL of methanol was added to each mixture and stirred, and the resultant mixtures were centrifuged. A total of 100 μL of the supernatant was collected from each mixture. Then, 150 μL of water was added, and the resultant solutions after stirring were used as the measurement samples. Chlorpromazine- containing plasma samples were purified by deproteinization. Subsequently, 10 μL of methanol and the IS solution 10 µL (0.05 μg/mL) were added to 20 μL of each plasma sample and stirred. Next, 100 μL of the borate pH standard solution (pH 9.18) and 700 μL of diethyl ether were added to each mixture and stirred, and the resultant mixtures were centrifuged. The organic solvent layer (700 μL) was collected from each mixture and dried under nitrogen at 40 degrees Celsius. One-hundred microliters of methanol was added to each residue and stirred. Then, 250 μL of 10 mmol/L ammonium formate solution/formic acid (1000:1 (v/v)) was added, and the resultant solutions after stirring were used as measurement samples.

### Instruments

High-performance liquid chromatography (HPLC) was performed using the Nexera system or LC-20A system (Shimadzu, Kyoto, Japan). The mass spectrometer was performed using API5000 (AB Sciex Pte. Ltd., Framingham, MA, USA). For MS control and spectral processing, Analyst software 1.6.1 (AB Sciex) was used.

### Liquid Chromatographic and Mass Spectrometric Conditions

SUMIPAX ODS Z-CLUE (50 mm × 2.0 mm i.d., 3 μm) (Sumika Chemical Analysis Service, Osaka, Japan) was used as the analytical column. For the analysis of azasetron, and domperidone, 10 mmol/L ammonium formate (pH 3.0) was used for mobile phase A and methanol was used for mobile phase B. Olanzapine were analyzed using mobile phase A with 10 mmol/L ammonium formate and mobile phase B with methanol. For the analysis of chlorpromazine, a mixture of 10 mmol/L ammonium formate solution and formic acid (1000:1 (v/v)) was used for mobile phase A and acetonitrile was used for mobile phase B. Two or ten microliter of each sample was injected into each column, and separation and elution were performed under the following gradient conditions: for azasetron, flow rate (mL/min): 0.3, mobile phase B (%): from 20–90; for olanzapine, flow rate (mL/min): 0.35, mobile phase B (%): from 35–95; for domperidone, flow rate (mL/min): 0.3, mobile phase B (%): from 30–90; and for chlorpromazine, flow rate (mL/min): 0.4, mobile phase B (%): 30–95. The column temperature was maintained at 40 degrees Celsius. Ionization was measured in Electrospray Ionization (ESI) mode and quantified by multiple reaction monitoring (MRM). The measurement conditions of MRM for each compound were as follows: azasetron, 350–224 m/z; azasetron IS, m/z 355–110 m/z; olanzapine, 313–256 m/z; olanzapine IS, 358–198 m/z; domperidone, 426–147 m/z; domperidone IS, 432–181 m/z; chlorpromazine, 319–86 m/z; and chlorpromazine IS, 325–92 m/z. The concentration ranges of each calibration curve were 2–2,000 ng/mL for azasetron and olanzapine, 0.2–200 ng/mL for domperidone, and 0.3–300 ng/mL for chlorpromazine.

### Data Analysis

Pharmacokinetic parameters were analyzed by a non-compartment model using Phoenix WinNonlin 6.3 (Pharsight Corporation, St. Louis, MO, USA), a pharmacokinetic analysis software. The following parameters were calculated from the plasma concentration-time data of each rat: maximum plasma concentration (*C*_max_), *T*_max_, elimination half-life (*t*_1/2_), elimination rate constant (*k*_el_), Area under the plasma concentration-time curve (AUC) from the time of administration to infinity (AUC_0-inf_), volume of distribution (Vd), and apparent clearance (CL). Statistical analysis was performed using a statistical analysis software GraphPad Prism 9 (Version 9.5.0, GraphPad Software, San Diego, CA, USA). Student’s t-test was used, and the significance level was set at *p* < 0.05.

## Results

### Skin thickness of SD rats and ID administration

The skin thickness of SD rats used in the present study was thinner than previously reported (1.18 mm) and was thinner than that of 8-week-old Wistar rats (average ± standard deviation; 0.60 ± 0.04 mm), but the success rate of ID administration with IDS for rats was 100% (Figure 2, Table 2) (30, 31).

**Figure 2.**
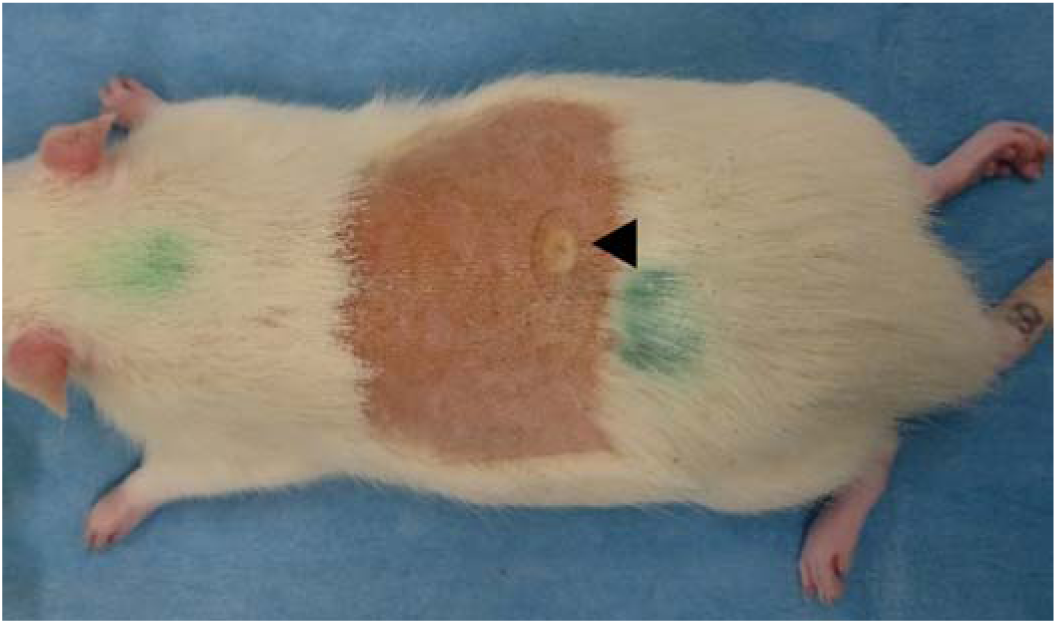
Wheal of ID administration (the representative sample). Arrow shows the wheal of ID administration.

**Table 2.** The Success rate of ID administration using Immucise for rats (n=16).

| Skin thickness (mm) | Criteria | Rates (%) |
| --- | --- | --- |
| Ave. 0.52 | Leakage at the injection site | 100 |
| S.D. 0.05 |  |  |
| Max. 0.60 | Wheal formation (> 3 mm) | 100 |
| Min. 0.44 | Successful ID injection<sup>a</sup> | 100 |

<sup>a</sup> Successful ID injection was defined as no visible leakage of the fluid on the epidermis or around the needle and wheal diameter of 3 mm or more.
Ave., Average; S.D., Standard deviation; Max., Maximum; Min., Minimum.

### Pharmacokinetics

The pharmacokinetic profiles of each administration route are shown in Figure 3, Table 3, and Supplemental figure.

**Figure 3.**
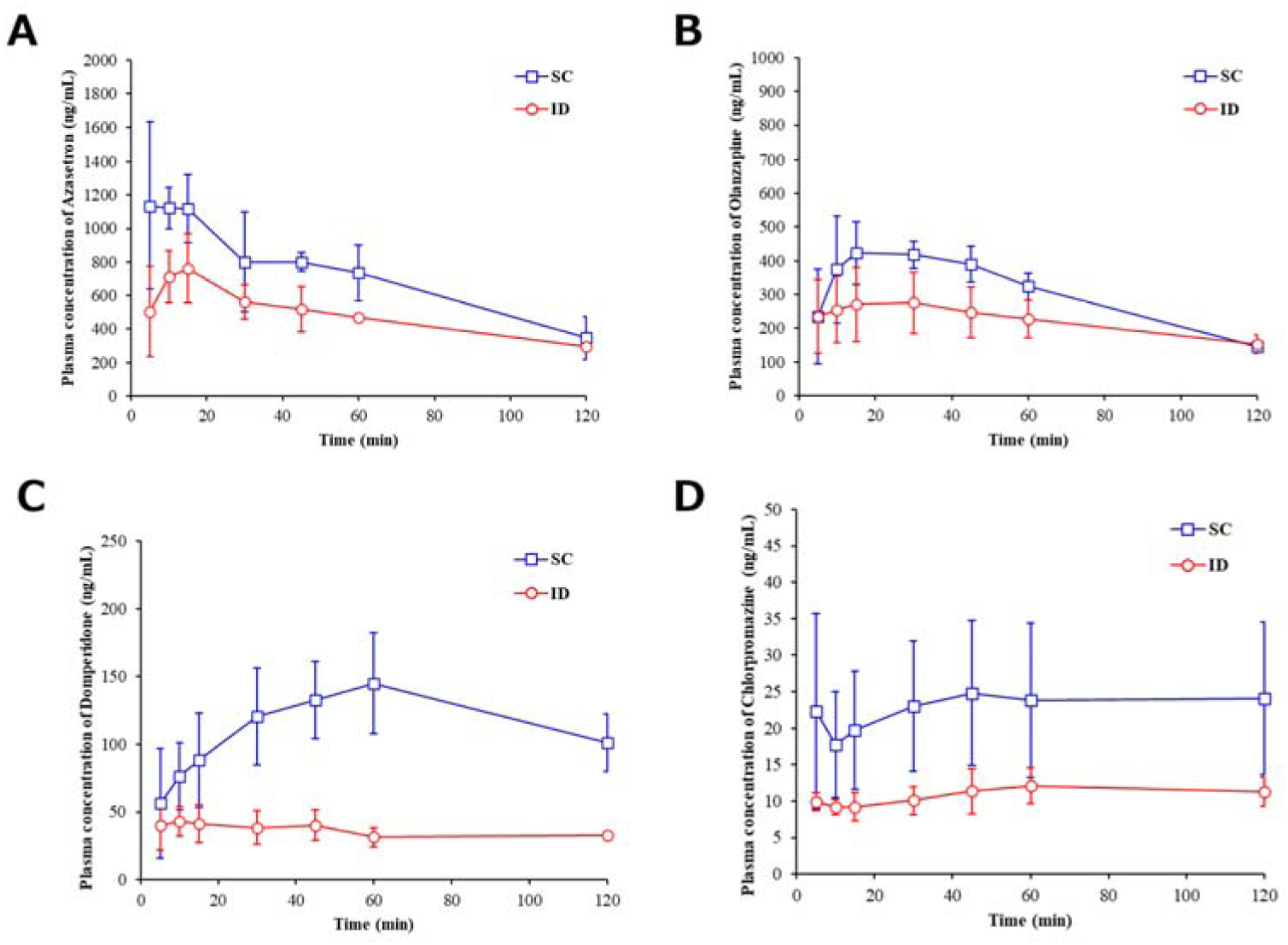
Mean plasma concentration-time profile from time zero to 120 minutes after administration. Mean plasma concentration-time profile from time zero to 120 minutes after single-dose SC or ID administration of azasetron (A), olanzapine (B), domperidone (C) or chlorpromazine (D) in rats. The error bars mean standard deviations. SC: subcutaneous, ID: intradermal.

**Table 3.** Plasma pharmacokinetics parameters after single-dose SC or ID administration of azasetron, olanzapin, domperidone, or chlorpromazine in rats.

| Parameter | SC | ID |
| --- | --- | --- |
| **Azasetron (n=3)** |  |  |
| $C_{\max}$ (ng/mL) | 1330 ± 340 | 816 ± 151 |
| $T_{\max}$ (min) | 11 ± 6 | 18 ± 10 |
| $t_{1/2}$ (hr) | 1.01 ± 0.30 | 1.80 ± 0.88 |
| $k_{el}$ (hr<sup>-1</sup>) | 0.730 ± 0.223 | 0.442 ± 0.171 |
| AUC<sub>0-inf</sub> (μg · hr/L) | 1950 ± 390 | 1710 ± 330 |
| Vd (L) | 1.38 ± 0.17 | 2.73 ± 0.81 * |
| CL (L/hr) | 0.989 ± 0.199 | 1.11 ± 0.18 |
| **Olanzapine (n=3)** |  |  |
| $C_{\max}$ (ng/mL) | 452 ± 88 | 282 ± 97 |
| $T_{\max}$ (min) | 28 ± 18 | 26 ± 9 |
| $t_{1/2}$ (hr) | 0.880 ± 0.013 | 1.95 ± 0.73 |
| $k_{el}$ (hr<sup>-1</sup>) | 0.788 ± 0.012 | 0.384 ± 0.118 * |
| AUC<sub>0-inf</sub> (µg·hr/L) | 779 ± 70 | 847 ± 60 |
| Vd (L) | 1.26 ± 0.10 | 2.55 ± 1.16 |
| CL (L/hr) | 0.988 ± 0.083 | 0.889 ± 0.076 |
| **Domperidone (n=5)** |  |  |
| C<sub>max</sub> (ng/mL) | 146 ± 37 | 49.5 ± 15.7 * |
| T<sub>max</sub> (min) | 58 ± 7 | 17 ± 16 |
| t<sub>1/2</sub> (hr) | 2.66 ± 1.40 | 4.71 ± 0.67 * |
| k<sub>el</sub> (hr<sup>-1</sup>) | 0.348 ± 0.214 | 0.150 ± 0.022 |
| AUC<sub>0-inf</sub> (µg·hr/L) | 619 ± 218 | 347 ± 41 * |
| Vd (L) | 2.78 ± 1.01 | 8.71 ± 1.83 * |
| CL (L/hr) | 0.828 ± 0.281 | 1.27 ± 0.15 * |
| **Chlorpromazine (n=5)** |  |  |
| C<sub>max</sub> (ng/mL) | 29.9 ± 9.8 | 12.8 ± 2.6 * |
| T<sub>max</sub> (min) | 118 ± 141 | 82 ± 91 |
| t<sub>1/2</sub> (hr) | 4.45 ± 1.96 | 11.2 ± 0.4 * |
| k<sub>el</sub> (hr<sup>-1</sup>) | 0.180 ± 0.071 | 0.0620 ± 0.0023 * |
| AUC<sub>0-inf</sub> (µg·hr/L) | 298 ± 77 | 202 ± 20 * |
| Vd (L) | 8.69 ± 3.17 | 31.6 ± 5.4 * |
| CL (L/hr) | 1.42 ± 0.37 | 1.95 ± 0.27 * |

Values are expressed as mean ± standard deviation. SC: subcutaneous, ID: intradermal, \*: Values of ID administration group were significantly different from those of SC administration group (p < 0.05).

SC administration in rats had a higher *C*_max_ than ID administration for all drugs, and had a significantly higher *C*_max_ for domperidone and chlorpromazine (domperidone: *p* < 0.001, chlorpromazine: *p* = 0.006). *T*_max_ after SC administration was fastest in chlorpromazine, followed by domperidone, olanzapine, and slowest in azasetron. *T*_max_ values of azasetron, olanzapine, and domperidone after ID administration were similar, but ID administration of domperidone showed a significantly faster *T*_max_ than SC administration (*p* < 0.001). ID administration showed longer *t*_1/2_ of all drugs compared with SC administration. Furthermore, *t*_1/2_ of domperidone and chlorpromazine were significantly longer (domperidone: *p* = 0.018, chlorpromazine: *p* < 0.001). ID administration of olanzapine and chlorpromazine showed a lower *k*_el_ than with SC administration (olanzapine: *p* = 0.004, chlorpromazine: *p* = 0.006). SC administration of domperidone and chlorpromazine showed significantly higher AUC_0-inf_ values than with ID administration (domperidone: *p* = 0.025; chlorpromazine: *p* = 0.027). ID administration of domperidone and chlorpromazine resulted in higher Vd and CL values than with SC administration (Vd: azasetron, *p* = 0.048; domperidone, *p* < 0.001; chlorpromazine, *p* < 0.001; CL: domperidone, *p* = 0.014; chlorpromazine, *p* = 0.032).

## Discussion

The successful ID administration rate showed that IDS for rats was suitable for ID administration not only in Wistar rats (30) but also in SD rats.

*T*_max_ values in single-dose PO administration to rats were reported to be 0.5–0.6 h for azasetron, 1 h for olanzapine, 2 h for domperidone, and 2–4 h for chlorpromazine, and the present study showed ID and SC administrations had lower *T*_max_ values for all these drugs (32–35). The blood profiles of chlorpromazine showed double peaks after ID and SC administration, and majority of the 1^st^ peaks were observed 5 min after dosing via either route (Figure 3). Chlorpromazine is known to easily enter enterohepatic circulation. Therefore, it was considered to be absorbed rapidly (the 1st peak) and then reabsorbed through the enterohepatic circulation (the 2^nd^ peak) (36). The concentrations of the 1^st^ and 2^nd^ peaks were similar, suggesting that the drug was rapidly transferred into the blood at the 1^st^ peak.

The resultant values of *C*_max_ and AUC_0-inf_ indicated that ID administration tended to show lower bioavailability than SC administration. Drugs administered by injection are generally more bioavailable than the same dose administered orally. Both ID and SC administrations may show faster *T*_max_ and higher bioavailability than PO, and these administration routes may be superior to PO in terms of onset of action and efficacy. As the effective blood concentrations of antiemetics in humans are unknown, further investigation is warranted.

The subcutaneously administered drug passes diffusively through the SC interstitium and reaches the capillaries or enters lymphatic vessels, which have large intercellular spaces and lack the basement membrane. Because small molecules and peptides pass through tight junctions present in capillaries and enter the blood, absorption uptake by lymph vessels is not predominant. In contrast, highly hydrophobic and high-molecular-weight drugs tend to be taken up by lymphatic vessels. It is known that drugs are absorbed slowly and their half-lives are relatively longer because the lymphatic flow rate is slow (37, 38). As the drug hydrophobicity is increased, *T*_max_ values after single SC doses of small-molecule drugs to rats increased as follows: within 0.08 h for radiolabeled ^14^C- sumatriptan succinate (sumatriptan’s logP = 0.93), 0.117 ± 0.017, 0.167 ± 0.017, 0.250 ± 0 h for ondansetron (logP = 2.40), and 0.5 h for risperidone (logP = 3.27) (39–41). A similar trend was observed in the results of SC administration in the present study. Contrarily, the dermis has a microcapillary bed composed of valveless blood vessels and open-ended lymph vessels. The intradermally administered drug is thought to enter the systemic circulation rapidly by being taken up by capillaries or lymph vessels (42). An earlier study reported a significantly rapid uptake of peptides and proteins into the blood after ID administration than after SC administration to pigs (43). The present study showed that domperidone was rapidly absorbed into the blood. The results also suggested that highly hydrophobic drugs such as domperidone are likely to be rapidly absorbed in blood by ID administration as opposed to SC administration.

Prior studies of antiemetic in humans have showed the potential of SC administration as an alternative for patients receiving platinum-based chemotherapy who cannot tolerate PO or IV administration (43, 44). SC injection devices including pre-filled syringes (PFS) and autoinjectors (AI) are universally used, and many products are indicated for self-injection. It has been reported that patients with rheumatoid arthritis prefer easy-to-use, convenient, less painful, and less-time consuming devices as self-injection devices that they can use at home (46). Auto-injectors (AIs), commonly used as SC devices for self-administration, are hand-grip and vertical-puncture type devices that are different from devices for normal SC injection and that improve patient compliance and adherence (47). However, many patients and their caregivers have a fear of needles, and this discomfort can delay or result in avoidance of medication (48). Selecting smaller needle sizes (both gauge and length) can reduce needle fear and pain, and increase injection comfort (18–21). Thus, it is expected to develop a highly effective and comfortable means of administration that can be used at home. Therefore, Immucise, which is a hand-grip and vertical-puncture type device similar to AIs, may be applicable as a self-administration device to be used at home. Also, compared to common SC needles, which are 1/2 inch (about 12 mm) to 7/8 inch (about 22 mm) long and their size are 23 to 25 gauge (49), Immucise has a shorter needle length and smaller needle diameter than general SC injection needles, which is expected to reduce pain (48, 50).

Immucise (Terumo, Tokyo, Japan) is an ID injection system that has been used in more than 750 patients in clinical studies has shown high rate of successful ID injection (51, 52). It has two types of syringes; prefilled and disposable, and the needles can be connected to both syringes (Figure 1A and 1 B). The prefillable syringe type Immucise was approved in Japan in 2015. The 510 (k) submission for the disposable type was approved by the FDA in 2018, and this obtained the Conformité Européenne (CE) mark in the European Economic Area in 2019. The Immucise structure, especially the needle length, is designed to be suitable for use in the human deltoid region. The skin thickness of the abdomen and thighs, which are generally used for self-injection, is slightly thinner than that of the deltoid region. (mean skin thickness of deltoid: 2.02 mm; mean skin thickness of abdomen: 1.91 mm; mean skin thickness of thigh: 1.55 mm) (53). Improving the structure of the needle, as in the modification in IDS for rats, could lead to the development of ID injection devices suitable for self-injection (30). Although it needs to be further verification such as Human Factors Engineering test by potential self-administration patients to ensure that it can be used comfortably at home, the improved needle of the Immucise could be used as the first ID self-administration device for liquid drug.

In this study, SC and ID administration of antiemetic showed the possibility of alternative route for the existing administration route. Also, an ID injection device with a smaller needle would be more effective than the typical SC administration device in terms of usability and pain reduction. Thus, the future application of vertical puncture administration devices such as Immucise for self-injection will provide simple and minimally invasive administration for rescue administration such as for breakthrough CINV and other events requiring rescue treatment at home.

## Conclusion

Single ID and SC administration of antiemetics to rats resulted in lower *T*_max_ values compared to previously reported oral administration. The results of this study suggest that ID and SC are useful routes of administration of antiemetics for breakthrough CINV at home and that the application of Immucise may provide an easy, ID-administered, home self-injection rescue antiemetic therapy.

## Abbreviations

AUC: Area under the plasma concentration-time curve
AUC_0-inf_: Area under the concentration-time curve from time zero to infinity
CE: Conformité Européenne
CINV: Chemotherapy-induced nausea and vomiting
CL: Apparent clearance
*C*_max_: Maximum plasma concentration after drug administration
ESMO: European Society for Medical Oncology
FDA: Food and Drug Administration
HEC: Highly emetogenic chemotherapy
HPLC: High Performance Liquid Chromatography
ID: Intradermal
IM: Intramuscular
Immucise: Immucise Intradermal Injection System
IS: Internal standard
IV: Intravenous
*k*_el_: Elimination rate constant
MASCC: Multinational Association of Supportive Care in Cancer
MEC: Moderately emetogenic chemotherapy
PO: Per os
SC: Subcutaneous
SD: Sprague-Dawley
*T*_max_: Time to maximum plasma concentration after drug administration
*t*_1/2_: Half-life time
Vd: Volume of distribution

## Author contributions

Minami Maekawa-Matsuura: Conceptualization, Methodology, Investigation, Writing - Original Draft, Project administration. Natsuko Shimizu: Investigation, Project administration. Yuri Sakamaki: Formal analysis, Investigation. Eri Oya: Investigation. Sakiko Shimizu: Writing - Review & Editing. Yoichiro Iwase: Supervision.

## Acknowledgements

The authors would like to thank members of Evaluation Center and Pharmaceutical Solutions Division of TERUMO CORPORATION for the assistance with the evaluations.

---

**Supplemental figure.**
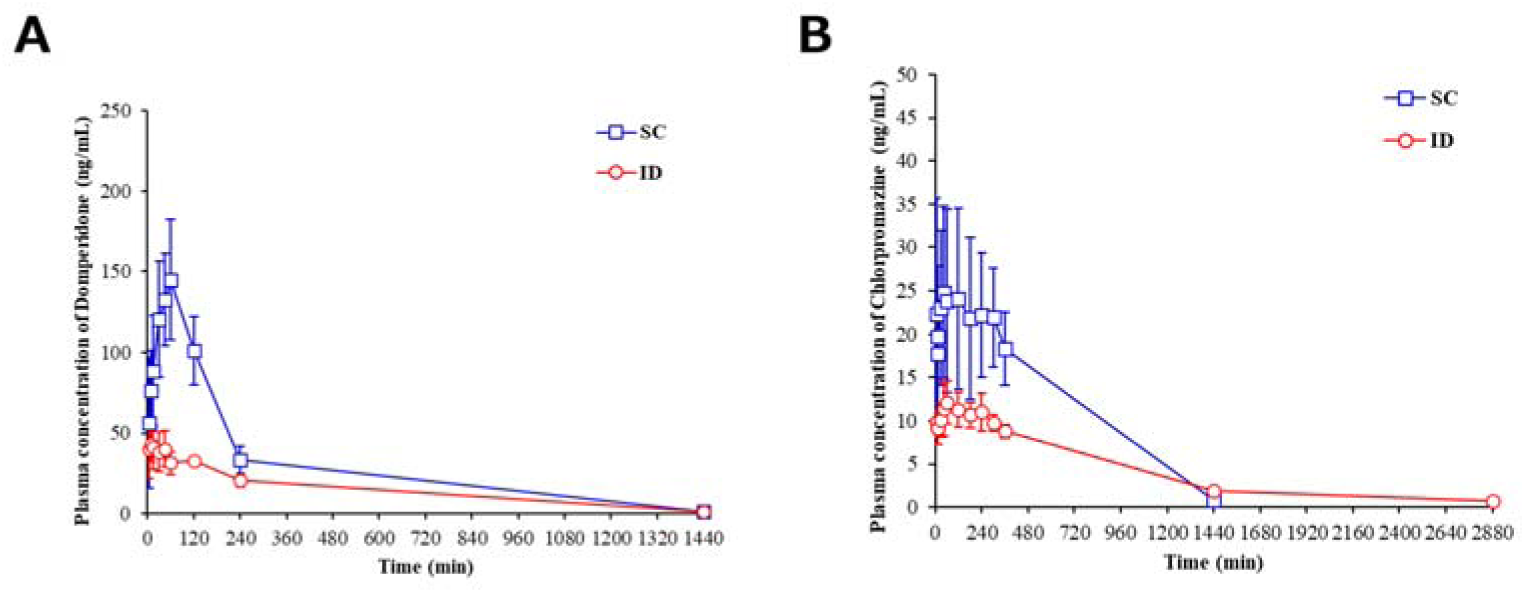
Mean plasma concentration-time profile after single-dose SC or ID administration of domperidone or chlorpromazine in rats. Mean plasma concentration-time profile after single-dose SC or ID administration of (A) domperidone 1.4 mg/kg or (B) chlorpromazine 1.2 mg/kg in Sprague-Dawley rats. The error bars mean standard deviations. SC: subcutaneous, ID: intradermal.

